# Plant MutS Homolog 1 is a mismatch-directed nuclease required for organelle genome maintenance

**DOI:** 10.64898/2026.06.11.731605

**Authors:** Alejandro Peñafiel-Ayala, Chang Zhou, Noe Baruch-Torres, Daniel B. Sloan, Shin-ichi Arimura, Luis G. Brieba

## Abstract

The exceptionally low mutation rates of plant organellar genomes imply the existence of DNA surveillance mechanisms that counteract replication errors and DNA damage. Genetic evidence implicates MutS HOMOLOG 1 (MSH1) as a central component of this pathway, as loss of MSH1 results in the accumulation of point mutations. MSH1 is a unique protein that combines an N-terminal MutS-like mismatch-recognition module with a C-terminal GIY–YIG nuclease domain. Here, we show that *Arabidopsis thaliana* MSH1 (AtMsh1) recognizes mismatches, insertion/deletion loops, and damaged bases within double-stranded DNA and introduces staggered DNA breaks at positions flanking the mismatch or lesion. Given the presence of an active homologous recombination machinery in plant organelles, we hypothesize that these DNA ends may be processed by exonucleases to remove the mismatched or damaged DNA while generating 3′ single-stranded DNA substrates suitable for homologous recombination-mediated repair and gene conversion. Together, our findings support a model in which AtMsh1 functions as a minimal mismatch repair system that couples mismatch recognition to DNA incision, providing a potential mechanism for suppressing mutation accumulation and maintaining the remarkable stability of plant organellar genomes.

**Key Points:**

- AtMsh1 recognizes and cleaves DNA mismatches, short insertion/deletion loops, and damaged DNA.
- AtMsh1 functions as a minimal mismatch repair system by coupling mismatch and lesion recognition to double-stranded DNA cleavage.
- The staggered 3′ single-stranded DNA ends generated by AtMsh1 cleavage are substrates for homologous recombination-mediated repair and gene conversion.

**Graphical Abstract:** Mismatch recognition and nuclease activity by plant organellar MutS Homolog 1 drive organellar genome maintenance.

## INTRODUCTION

In flowering plants, mitochondrial genomes exhibit exceptionally low synonymous substitution rates, approximately 100-fold lower than those of animal mitochondria and 16-fold lower than in plant nuclear genomes. Plastid genomes also evolve slowly, with synonymous substitution rates averaging approximately five-fold lower than those of the nuclear genome ^1,2^. These low synonymous substitution rates are particularly striking given the relatively low fidelity of plant organellar DNA polymerases and the chemical milieu that damages DNA in mitochondria and plastids. These observations imply the existence of highly efficient DNA surveillance and repair mechanisms that compensate for replication errors and DNA damage, thereby preserving organellar genome integrity ^1–4^. *MutS HOMOLOG 1* (*MSH1*) is necessary for maintaining the exceptionally low mutation rates of mitochondrial and plastid DNA while suppressing ectopic recombination and promoting organellar homoplasmy ^5–9^. Sequence homology suggests that Msh1 is a bifunctional protein composed of an N-terminal MutS1 DNA mismatch repair (MMR) module fused to a C-terminal GIY-YIG nuclease domain ^5,6,8,10–14^. The N-terminal MutS1 module of MSH1 is homologous to bacterial MutS1 proteins, the ATP-dependent mismatch sensors that initiate MMR. During bacterial MMR, MutS1 functions as a homodimer that scans duplex DNA for the helical distortions generated by mismatched base pairs and insertions/deletions (indels). Upon mismatch recognition, MutS1 recruits downstream repair factors that cleave the nascent DNA strand containing the replication error, followed by DNA resynthesis to restore the correct sequence ^15,16^. Although Msh1 retains the conserved mismatch-recognition and ATPase domains of MutS1, other components of the bacterial MMR machinery are absent in plant organelles. The presence of a GIY-YIG nuclease domain in MSH1 raises the possibility that mismatch recognition and DNA incision are directly coupled within a single polypeptide. The C-terminal GIY-YIG domain of Msh1 belongs to a widespread family of metal-dependent endonucleases found in restriction enzymes, homing endonucleases, structure-specific nucleases, and DNA repair proteins. GIY-YIG nucleases adopt an α/β-sandwich fold in which catalysis is mediated within the GIY and YIG motifs and conserved Arg, His/Tyr, and Glu residues that coordinate a catalytic metal ion ^17,18^.

MSH1 is found in green plants, some other plastid-containing eukaryotes, and giant viruses (nucleocytoviruses) but is absent in most other groups, including animals ^5,10,19,20^. Arabidopsis mutant lines devoid of functional Msh1 (AtMsh1) exhibit elevated rates of structural rearrangements, indels, and nucleotide substitutions in mitochondrial and plastid genomes ^5,7,11^. The fact that Arabidopsis *msh1* mutants exhibit mutation patterns similar to those observed in bacteria lacking MutS1 leads us to hypothesize that AtMsh1 may recognize mismatches and damaged DNA within the double-stranded DNA helix ^21–25^. Here, we present a comprehensive analysis of AtMsh1 recombinantly expressed in yeast, showing that this enzyme is a minimal mismatch repair (MMR) system capable of both recognizing and cleaving mismatches, leading to the formation of double-stranded breaks (DSBs). These findings uncover an effective MMR pathway in plant organelles and suggest that AtMsh1 provides surveillance for replication errors and DNA lesions.

## METHODS

### Subcloning of full-length AtMsh1 and site-directed mutagenesis

A synthetic construct containing the codon-optimized nucleotide sequence for AtMsh1 was designed for expression in both bacteria and yeast (Biomatik, Wilmington, DE, USA). The AtMsh1 (AT3G24320) coding region, excluding the predicted organellar targeting transit peptide (residues 1 to 31), was PCR-amplified using a forward primer containing a *Bam*HI site that introduced an initial methionine (Met), followed by a serine (Ser) and a stretch of ten histidine residues, positioned just upstream of the DNA sequence corresponding to AtMsh1 Pro32. The reverse primer was complementary to the region ending at Leu1118, including the stop codon and a *Xho*I restriction site. The sequences of the primers used for amplification are provided in **Table S1**. The resulting PCR product was cloned into the pSc-ADAR plasmid to generate the pSc-AtMsh1 construct. The pSc-ADAR plasmid is derived from YEpTOP2PGAL, contains a pBR322 backbone, and carries a URA3 selection marker, a 2µ origin of replication, an ampicillin resistance gene, and a GAL1 promoter for galactose-inducible expression ^26^. Point mutants were constructed following the Q5 Site directed mutagenesis protocol from New England Biolabs using specific oligonucleotides for Glu1078Ala and for Phe147Ala mutations as indicated in **Table S1**.

### Heterologous expression and purification of yeast recombinant wild-type and mutant AtMsh1

Yeast cells (BCY123) were cultured, treated with lithium acetate solution, and combined with pSc-AtMSH1, boiled salmon sperm DNA, and a 40% PEG solution to promote transformation. Following incubation at 30 °C and a 42 °C heat shock, cells were plated onto selective media and incubated at 30 °C for 2–3 days to allow growth of transformants. Overnight culture in YP + 2% glucose was used to inoculate 4L of YP + 0.5% glucose at 30 °C 300 rpm for 18 hours and then diluted to an O.D. value of 1.8 with fresh YP media. The culture was induced for 16 hours with a final concentration of 3% D-galactose. Cells were harvested after induction and resuspended in lysis buffer (25 mM HEPES-KOH 7.5, 1M NaCl, 5 mM imidazole, 10% glycerol, 1 mM PMSF), lysed, and centrifuged at 15,000 rpm for 60 minutes. Clear lysate was loaded into a HisTrap resin and washed with lysis buffer supplemented with 40 mM imidazole. Protein was eluted with lysis buffer supplemented with 500 mM imidazole. The eluate was loaded into a HP-Heparin column and washed with an increasing gradient of NaCl in buffer B (25 mM HEPES-KOH 7.5, 10% glycerol, 1 mM DTT, 1 mM PMSF). Fractions containing AtMSH1 were applied to an S-300 column pre-equilibrated with buffer D (25 mM HEPES-KOH 7.5, 300 mM NaCl, 1 mM DTT, 1 mM PMSF, 1% glycerol), using an ÄKTA pure system. Protein fractions were concentrated and stored at -80 °C. All proteins were snap frozen with liquid nitrogen.

### Structural Modeling

An AlphaFold 3 model of AtMsh1 was built using the elements from the crystal structure of *Thermus aquaticus* MutS1 bound to a G:T mismatch-containing DNA duplex (PDB: 1E3M) as a template ^22,27^. The AtMsh1 model preserved the DNA duplex, including the G:T mismatch, and the catalytic Mg²⁺ ion from the template, while replacing the bound ADP with ATP to represent the ATP-bound state of AtMsh1. Model quality was assessed using the predicted Local Distance Difference Test (pLDDT) scores generated by AlphaFold 3.

### Determination of the oligomeric status of AtMsh1

Size exclusion chromatography was performed on an ÄKTA-FPLC system using a Superose 12 10/300 GL column. The column was equilibrated with buffer containing 25 mM HEPES-KOH (pH 7.5), 5% glycerol, 300 mM NaCl, and 1 mM DTT. AtMSH1 was injected at a concentration of 2 mg/mL and eluted at a flow rate of 0.3 mL/min. Elution was monitored by absorbance at 280 nm. Gel filtration standards (Bio-Rad), including vitamin B12, myoglobin, ovoalbumin, gamma globulin, and thyroglobulin, were used to generate a molecular weight calibration curve.

### Electrophoretic Mobility Shift Assay (EMSA)

Fluorescently labeled oligonucleotides (**Table S2 and S3**) were quantified and annealed in 1x buffer H (10 mM Tris-HCl 7.5, 2 mM MgCl_2_, 100 mM NaCl) in an equimolar ratio. DNA oligonucleotides were annealed in a thermocycler using the following program: (I) 95 °C for 5 min (II) 95 °C – 1 °C/cycle (III) Goto 2, 70x, and (IV) and hold at 4 °C. To test for DNA binding interactions, 10 μL reaction volumes were set up using 2.5 nM DNA substrate containing reaction buffer without MgCl_2_ and with varying enzyme concentrations. Reactions were incubated for 25 minutes at room temperature, stopped using 8% glycerol, and loaded in a 10% polyacrylamide native 1x TBE gel. The gels were run in a 1x TBE buffer at 90 V for 2 hours and visualized using an Amersham Typhoon™ scanner (Cytiva) with Cy2 or Cy5 filters. Band intensities were quantified using ImageJ software. Data was analyzed using PRISM software and a one-site specific binding equation:

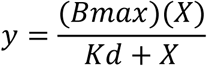

where X is the concentration of ligand, y is the specific binding and Kd is the equilibrium dissociation constant, and *Bmax* is the estimated plateau of the binding curve. Binding experiments were executed in triplicate.

### DNA cleavage reactions using synthetic oligonucleotides

Fluorescently labeled dsDNA substrates **(Table S3)** were incubated at 30 °C in a reaction buffer containing 25 mM BisTris propane (pH 7.5), 50 mM potassium glutamate, 5 mM MnCl₂ or MgCl₂, 1 mM DTT, 0.1% BSA, and 1 mM ATP, γ-ATP, or ADP. Reactions, typically containing 15 nM DNA substrate and varying protein concentrations, were stopped at different time points. Products separated by 12% denaturing PAGE in 1x TBE buffer were stopped with 2x stop buffer (95% formamide, 5 mM EDTA). Maxam-Gilbert reactions were performed with fluorescent oligonucleotides to precisely determine the cleavage site (80). 5 nM DNA substrate was separated on a native 12% polyacrylamide gel to preserve the dsDNA structure and stopped with 2x native stop buffer (8% glycerol, 0.6% SDS, 40 mM EDTA). Reactions were visualized using an Amersham Typhoon™ scanner (Cytiva) with Cy2 or Cy5 filters, and band intensities were quantified using ImageJ software.

## RESULTS

### AtMSH1 integrates the minimal functional elements required for mismatch repair within a single polypeptide

AtMsh1 is a member of a distinct subfamily of MutS1 proteins that possess a C-terminal GIY–YIG nuclease domain ^10^. Beyond the acquired nuclease module, AtMsh1 retains the conserved domain organization of bacterial MutS1, consisting of the canonical mismatch-binding (I), connector (II), core (III), clamp (IV), and ATPase (V) domains (**Fig. 1A-B**). An AlphaFold 3 structural model of AtMsh1 generated with the structural elements of *T. aquaticus* MutS1 bound to a T:G mismatch (PDB: 1E3M) predicts that AtMsh1 assembles as a homodimer through interactions between the ATPase domains of both monomers (**Fig. 1 A-B**) ^22^. Each GIY–YIG nuclease domain is connected to the C-terminal of the MutS1 domain by a flexible linker. Overall, the model exhibits high and continuous confidence scores across both domains, and more than 85% of the MutS1 domain residues have predicted Local Distance Difference Test (pLDDT) scores above 90 (**Fig. S1**). The GIY–YIG nuclease domain has an average pLDDT score of 70, and the regions of lower confidence are confined to the first 33 residues, residues 955–1004 corresponding to the linker connecting the MutS1 and GIY–YIG nuclease domains, and the final 19 residues of the nuclease domain. (**Fig. S1**). In order to biochemically characterize AtMsh1 and to overcome the low yields obtained during AtMsh1 purification expressed from bacteria ^14^, we expressed and purified recombinant AtMsh1 from yeast. Yeast-expressed purified AtMsh1 was >95% pure and eluted by size-exclusion chromatography with an apparent molecular weight consistent with a homodimer (**Fig. S2 A-D**). Given that mismatch-processing nucleases are sensitive to buffer composition ^28,29^, we screened AtMsh1 enzymatic activity using twelve reaction buffers (**Fig. S2 E**) that support nuclease activity of several enzymes. AtMsh1 cleaved the T:G mismatch-containing substrate under eight of the twelve conditions. Potassium was present in six of the eight buffer conditions that supported mismatch-directed nuclease activity, suggesting that, as in other DNA repair nucleases, this ion promotes enzymatic activity ^30–32^ (**Fig. S2 E**).

**Fig. 1.**
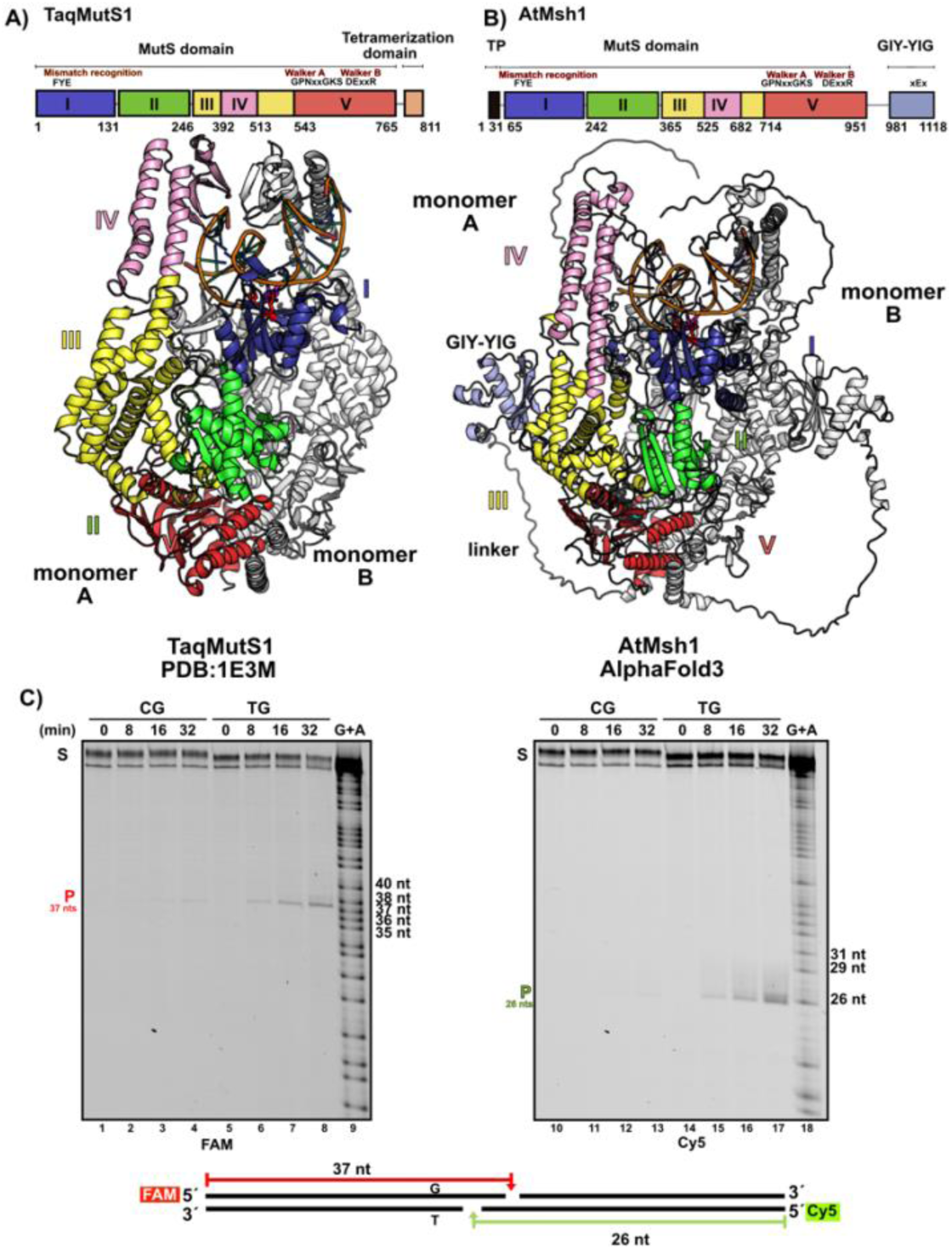
AtMsh1 is a minimal mismatch DNA repair system. A-B) Structural comparison of the AtMsh1 homology model and the crystal structure of *T. aquaticus* MutS1 bound to a mismatched DNA substrate (PDB: 1E3M). Both proteins are shown as homodimers, with the DNA-contacting monomer colored according to its domain organization and the second monomer shown in white. Duplex DNA is shown in orange. The nucleotide and catalytic metal ion bound to the ATPase domain (ADP and Mg²⁺ in *T. aquaticus* MutS1 and ATP and Mg²⁺ in AtMsh1) are represented as spheres and colored black and green, respectively. AtMsh1 exhibits the canonical MutS1 mismatch-binding (I), connector (II), core (III), clamp (IV), and ATPase (V) domains but differs from bacterial MutS1 proteins by the presence of a C-terminal GIY–YIG nuclease domain. **C)** 12% denaturing polyacrylamide gel showing the dsDNA cleavage of the mismatch-containing and complementary strands of a T:G mismatch in comparison to the absence of DNA cleavage on a non-mismatched DNA over a 32-min time course.

To further characterize AtMsh1, we used differential labeling of the DNA strands to discern strand-specific cleavage. Throughout this study, we refer to the thymine-containing strand from a T:G mismatch as the “error-containing” strand or “mismatch-containing” strand because studies of bacterial MutS1 proteins have shown it to specifically recognize the Thy base in T:G mismatches using stacking interactions via their mismatch-recognition motif ^22,33^. Conversely, we refer to the guanosine-containing strand as the “correct” or complementary strand. Using the optimized reaction conditions, we compared the nuclease activity of AtMsh1 on a dsDNA containing a single T:G mismatch and a fully base-paired dsDNA duplex over a 32 min time course (**Fig. 1C**). AtMsh1 efficiently cleaved both strands of the T:G mismatch substrate, whereas minimal cleavage of the canonical C:G duplex substrate was observed throughout the time course. Cleavage products from the T:G mismatch substrate increased with incubation time, indicating that AtMsh1 functions as a mismatch-directed dsDNA endonuclease (**Fig. 1C**). Detailed mapping of the AtMsh1 cleavage products using DNA sequencing reactions showed that AtMsh1 primarily cleaves the mismatch-containing strand 26 nts from its 5′ end (nine nts from the mismatch) and 37 nts from the 5′ end of the complementary strand (12 nts from the mismatch), generating 3′-staggered DNA ends with three-nt overhangs (**Fig. S3)**.

### AtMsh1 generates restriction-endonuclease-like staggered double-strand breaks at dsDNA mismatches and indels

As bacterial MutS1 proteins efficiently recognize indel loops of one and two nucleotides, we next examined the cleavage pattern of AtMsh1 using duplex substrates containing one or two thymidine insertions and compared them with a T:G mismatch substrate. AtMsh1 generated cleavage products on the insertion-containing strand that were one or two nucleotides longer than the predominant product produced from the T:G mismatch substrate, consistent with the presence of one or two additional thymidines (**Fig. 2A, B**). In contrast, cleavage of the complementary strand, which is identical in length in all three duplexes, produced the same predominant 26-nt product regardless of the substrate. These findings indicate that the GIY–YIG nuclease domain cleaves at a fixed distance from the boundary between distorted and canonical duplex DNA, using the first Watson–Crick base pair after the mismatch or indel as a positional landmark. This mechanism ensures that cleavage is phased relative to the mismatch or indel of this substrate, positioning the GIY–YIG active site nine nucleotides from the mismatch or indel on the mismatch-containing strand and 12 nucleotides from the mismatch or indel on the complementary strand, thereby generating 3′-staggered double-strand breaks with 3-nt overhangs (**Fig. 2A, B**).

**Fig. 2.**
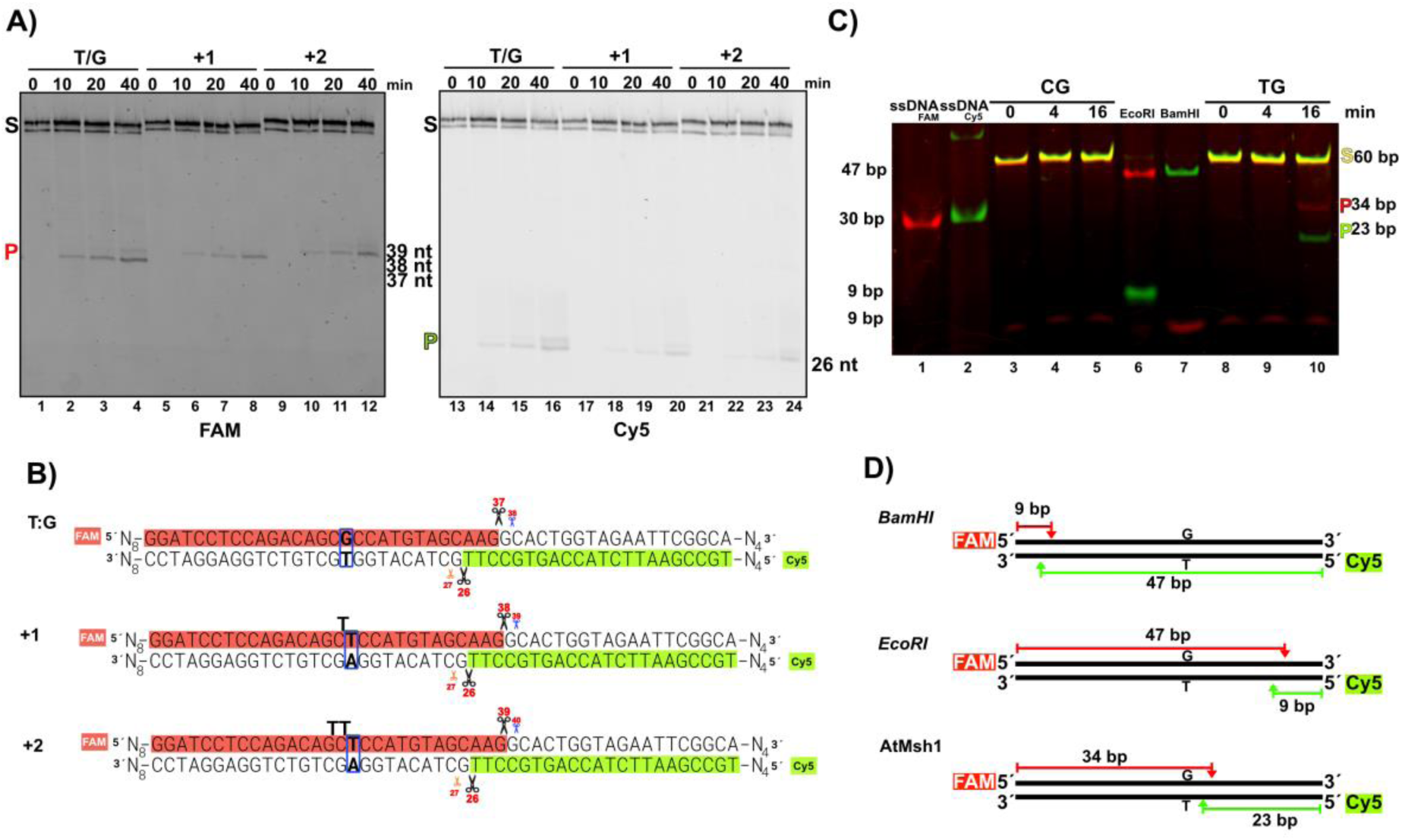
AtMsh1 cleaves mismatches and indel substrates and functions as a bona fide endonuclease. **A)** 12% denaturing polyacrylamide gel showing AtMsh1-mediated cleavage of DNA substrates containing a T:G mismatch or 1- and 2-nt insertions. The expected cleavage products of 37, 38, and 39 nts for the T:G mismatch, +1, and +2 insertion substrates, respectively, in the Gua-containing strand, and the corresponding 26-nt product in the mismatch-containing strand are indicated. **B)** Schematic representation of AtMsh1 cleavage products for mismatches and insertions of one and two nucleotides. The cleavage sites are indicated by scissors. **C)** Cleavage products generated by AtMsh1 resolved on native polyacrylamide gels to preserve native dsDNA structure. Control experiment showing double-stranded DNA cleavage products resulting from incubation with *Bam*HI or *Eco*RI, which have recognition sites that are located in opposite sides of the duplex DNA, as well as products following incubation with AtMsh1 on a G:T mismatch and a canonical G:C base pair. Non-annealed ssDNA controls were included to exclude the possibility that the observed cleavage products resulted from nonproductive annealing (lanes 1 and2). AtMsh1 cleavage generates a 5’-FAM-labeled 34-bp dsDNA fragment with a 3-nts 3′ overhang (red labeled), and a 5′-Cy5– labeled 23-nt dsDNA fragment, also bearing a 3-nt 3′ overhang (green labeled). **D)** Schematic representation of AtMsh1 cleavage showing the product lengths generated by AtMsh1 and the control restriction enzymes. Double-stranded DNA substrates containing *Bam*HI and *Eco*RI sites positioned on opposite sides of the duplex were used as controls to define the expected migration of the cleavage products.

To further substantiate that AtMsh1 acts as a mismatch-directed endonuclease, we performed dsDNA cleavage assays under native conditions using a dual-fluorophore–labeled substrate in comparison to DNA cleavage using restriction enzymes. Individual single-stranded DNA molecules labeled with Cy5 or FAM are visualized in red or green, respectively. Upon annealing, the resulting double-stranded DNA substrates display a yellow signal resulting from the overlap of the red and green fluorescence channels (**Fig. 2C, lanes 1, 2, 3, and 8**). Reactions incubated with *Eco*RI and *Bam*HI restriction enzymes generated fragments of nearly symmetrical length that carry opposite fluorophores and therefore display different colors (**Fig. 2C, lanes 6 and 7)**. In this experiment, Cy5-labeled and FAM-labeled dsDNA fragments of approximately 47 bp are visualized as green and red bands in reactions incubated with *Eco*RI and *Bam*HI, respectively (**Fig. 2D**). The 9-bp fragment generated by *Eco*RI cleavage exhibited anomalous migration, consistent with previously reported observations for Cy5-labeled DNA fragments of short length ^34^, in comparison to the migration of a 9-bp FAM-labeled dsDNA segment observed in a reaction incubated with *Bam*HI (**Fig. 2C, lanes 6 and 7**). In the presence of a mismatch, but not on canonical templates, AtMsh1 generated dsDNA cleavage products of approximately 34 and 23-nts, along with their corresponding overhangs, which precisely matched the fragment lengths observed under denaturing conditions (**Fig. 2C, lanes 4, 5, 9 and 10)** These results establish that AtMsh1 endonuclease activity generates DSBs in proximity to DNA mismatches (**Fig. 2C and 2D**). These results demonstrate that AtMsh1 cleaves both DNA strands to generate a site-specific dsDNA break at a fixed distance from the mismatch recognition site.

### The mismatch-recognition motif and ATPase activity are both required for AtMsh1-mediated cleavage of mismatch-containing DNA substrates

MutS1 proteins recognize mismatches through a universally conserved Phe-X-Glu motif, in which an invariant phenylalanine (Phe36 or Phe39 in *E. coli* and *T. aquaticus* MutS1, respectively) intercalates into the DNA duplex and directly contacts the mismatched base, while a conserved glutamate (Glu 41 in *T. aquaticus*) stabilizes the mismatch. Amino acid sequence alignment of plant Msh1 and bacterial MutS1 proteins reveals conservation of the mismatch-recognition motif across plants. Specifically, residues AtMsh1-Phe147 and AtMsh1-Glu149 occupy positions equivalent to the universally conserved Phe and Glu residues of bacterial MutS1 proteins, suggesting that plant Msh1 recognizes mismatched DNA through the same motif present in bacterial MutS1 enzymes (**Fig. 3A**). According to our structural model, AtMsh1 is predicted to recognize mismatched DNA following a canonical MutS1 mechanism. Residues AtMsh1-Phe147 and Glu149 superimpose with the Phe-X-Glu mismatch-recognition motif of bacterial MutS1 proteins, positioning residue AtMsh1-Phe147 to intercalate into the duplex from the minor groove and stack against the mismatched Thy, while AtMsh1-Glu149 is poised to stabilize the mismatched base through hydrogen bonding (**Fig. 3B**). To investigate this prediction, we constructed and purified an AtMsh1-Phe147Ala mutant enzyme **(Fig. S2C)**, as the corresponding mutation abolishes mismatch recognition in bacterial MutS1 ^35^. Consistent with our prediction, AtMsh1-Phe147Ala failed to cleave T:G-mismatched DNA in a side-by-side comparison with wild-type AtMsh1 (**Fig. 3C**). These results demonstrate that mismatch recognition is required to activate AtMsh1 nuclease activity and suggests that AtMsh1 integrates endonuclease activity in a compact MMR-like assembly ^36–40^.

**Fig. 3.**
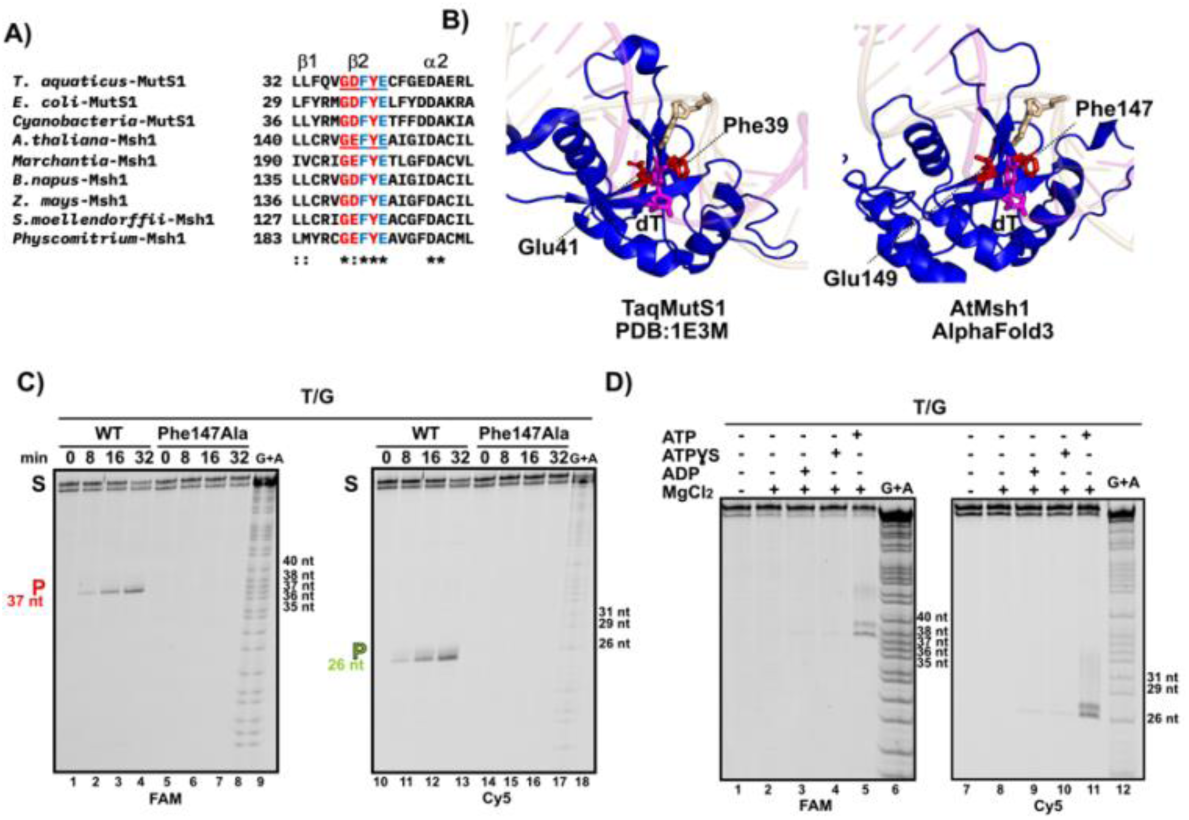
The conserved Phe147 residue and ATP hydrolysis are required for mismatch recognition and AtMsh1 nuclease activation. **A)** Amino acid sequence alignment of the mismatch-recognition motif of plant Msh1 enzymes in comparisons to mismatch-recognizing MutS1 enzymes from bacteria. The conserved mismatch-recognizing Phe–X–Glu motif is colored in cyan. The alignment illustrates the amino acid conservation across bacterial MutS1 proteins and Msh1 proteins from diverse Viridiplantae lineages. **B)** Ribbon representation of a structural comparison of the mismatch-recognition sites of *T. aquaticus* MutS1 bound to a T:G mismatch (PDB: 1E3M) and the AlphaFold 3 model of AtMsh1. The conserved Phe and Glu residues. (red colored) are positioned near the mismatched Thy (colored pink) whereas the Gua base is shown in wheat color **C)** AtMsh1-Phe147Ala is deficient in cleavage of T:G-mismatched DNA. Denaturing gel showing cleavage products from a T:G-containing DNA substrate in a side-by-side assay using wild-type AtMsh1 and AtMsh1-Phe147Ala reaction. DNA strands were labeled at the 5′ end with FAM (left) or Cy5 (right). The positions of the substrate (S) and cleavage products (P) are indicated. **D)** AtMsh1 is a mismatch-recognizing nuclease whose catalytic activity depends on ATP binding and a divalent metal ion. 12% denaturing polyacrylamide gel showing the endonucleolytic cleavage of the mismatch-containing and correct strands AtMsh1 nuclease activity requires ATP hydrolysis and a divalent metal ion. ADP and ATPγS support only weak cleavage, whereas ATP strongly stimulates cleavage and produces additional products on both DNA strands.

MutS1 and Msh1 enzymes belong to the ATP-binding cassette (ABC) transporter protein family ^41^. During mismatch recognition, bacterial MutS1 enzymes undergo ATP-driven conformational changes in which they transition from a mismatch recognition state into a sliding-clamp state that enables recruitment of the components of the mismatch-repair pathway ^15,16^. We examined whether ATP hydrolysis is required for DNA cleavage by comparing reactions containing ATP, the non-hydrolyzable analog ATPγS, and ADP (**Fig. 3D**). In the presence of ADP or ATPγS, AtMsh1 generates a faint cleavage product on both the mismatch-containing strand and the complementary strand; however, in the presence of ATP, these bands become much more pronounced, and an additional product 1 nt longer appears in both strands (**Fig. 3D**). These cleavage products are the only ones found in the presence of ADP or ATPγS and are detected at early time points in reactions incubated with ATP. In contrast, fainter products of 27 and 38 nts arise when incubated with ATP (**Fig. 3D, lanes 5 and 12**).

### The GIY–YIG nuclease active site is required for cleavage mismatch-containing DNA substrates

Amino acid sequence alignment of the GIY–YIG nuclease domain of AtMsh1 with Msh1 proteins from other eukaryotes and the well-characterized GIY–YIG nucleases I-TevI, Eco29kI, and UvrC, illustrates that the catalytic GIY, YIG, and RXXXH/Y motifs and the catalytically essential glutamate residue are conserved in Msh1 proteins (**Fig. 4A**). Structural modeling further indicates that these conserved residues in the AtMsh1 GIY–YIG domain are predicted to occupy positions corresponding to those of the catalytic residues in the homing endonuclease I-TevI, suggesting a conserved active-site architecture and a potential single-divalent-metal-ion mechanism of catalysis ^42,43^ (**Fig. 4B**). To determine whether the GIY–YIG nuclease domain mediates mismatch-directed DNA cleavage, we generated and purified a predicted catalytic-site mutant defective in DNA cleavage, in which the conserved AtMsh1-Glu1078 residue was substituted with alanine (Glu1078Ala) **(Fig. S2C)** and compared its cleavage activity with that of wild-type AtMsh1 using a dsDNA substrate containing a T:G mismatch. (**Fig. 4C**). The nuclease mutant failed to cleave the mismatched substrate under identical reaction conditions as wild-type AtMsh1, demonstrating that mismatch-directed DNA cleavage depends on the GIY– YIG domain (**Fig. 4C**).

**Fig. 4.**
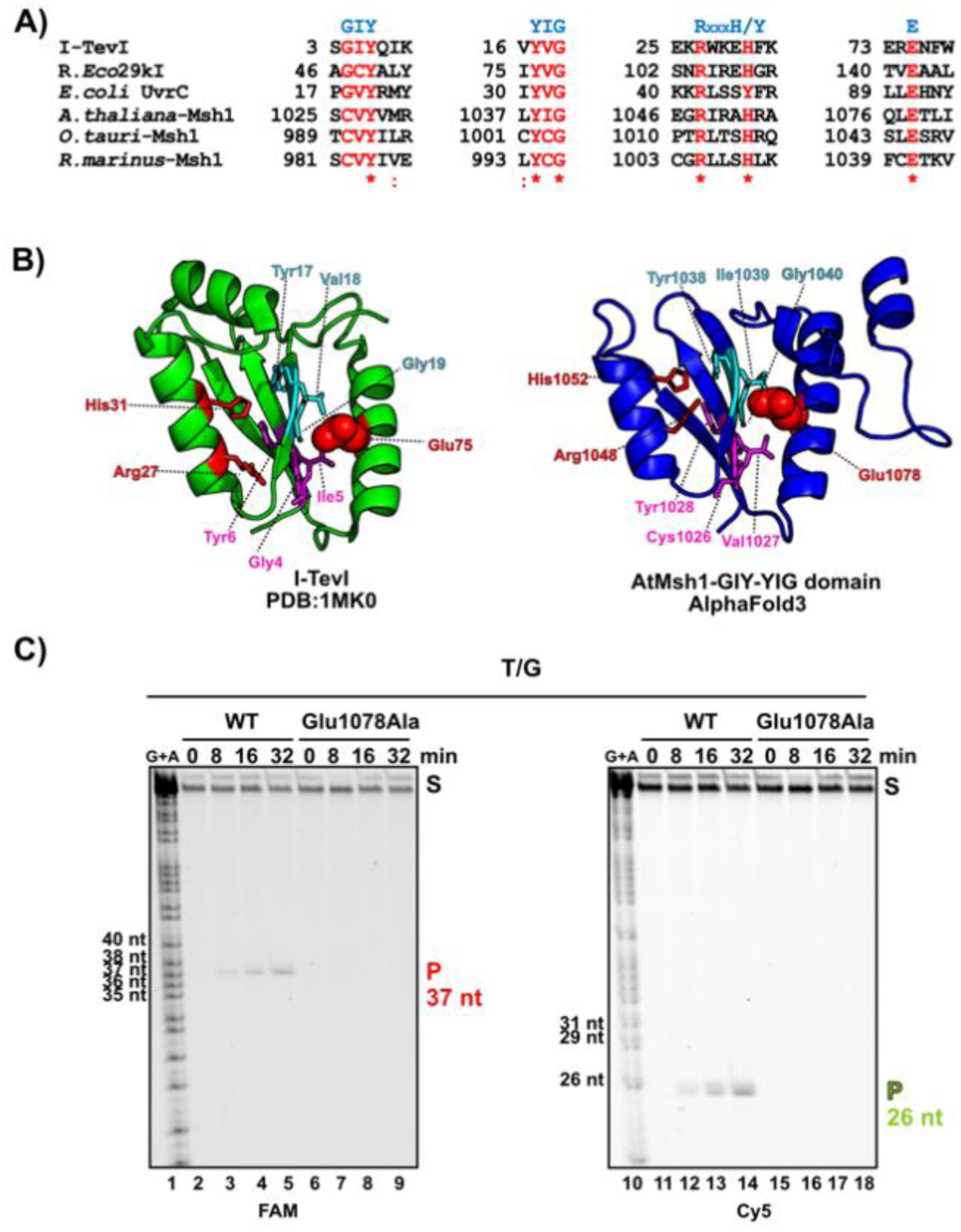
The GIY–YIG nuclease active site is required for cleavage mismatch-containing DNA substrates. **A)** Multiple sequence alignment of the catalytic motifs of representative GIY–YIG nucleases, with *A. thaliana*, *Ostreococcus tauri*, and *Rhodosorus marinus* Msh1. Conserved and catalytic residues are highlighted in red, showing the GIY, YIG, RxxxH/Y motifs and the invariant Glu**. B)** Structural model of the AtMsh1 GIY–YIG nuclease domain (residues 1022–1118), shown in blue and oriented to match the crystal structure of I-TevI ^42^ in green, illustrating conservation of the catalytic fold and active-site residues. Predicted catalytic residues are shown in ball-and-stick representation. The GIY and YIG domains are colored purple and cyan, respectively, the conserved RXXXH/Y motif is highlighted in red, and the conserved catalytic Glu residue is shown as spheres (I-TevI-Glu75 and AtMsh1-Glu1078Ala). **C)** A 12% denaturing polyacrylamide gel showing the loss of endonucleolytic cleavage activity by the AtMsh1-Glu1048Ala mutant compared with the dsDNA cleavage by wild-type AtMsh1.

### AtMsh1 exhibits distinct recognition and cleavage specificities for DNA mismatches and indels

Bacterial MutS1 preferentially binds 1-or 2-nt indels and G:T mismatches, whereas other mismatches (e.g., C:C, C:T, G:A, A:C) are recognized with affinities similar to duplex DNA ^44^. To assess whether AtMsh1 exhibits a substrate preference profile, we conducted electrophoretic mobility shift assays (EMSAs) to determine binding constants for all mismatches and indels (**Fig. 5A**). We found that AtMsh1 exhibits a preferred binding to indels of 1 and 2 nts followed by binding to T:G, T:T, and T:C mismatches (**Fig. 5A to D**). The binding affinities of AtMsh1 contrasts with MutS1 from *E. coli*, which displays highest affinity for G:T mismatches and ∼18-fold weaker binding to C:C mismatches ^45^. To assess whether substrate binding and dsDNA cleavage are correlated, we evaluated the substrate cleavage preference of AtMsh1 using increasing enzyme concentrations (0.25 to 1 nM) while maintaining a constant incubation time (**Fig. 5E to G**). Under those conditions several mismatches (specifically T:G, T:C, T:T, and A:C) were cleaved with high efficiency, a second class of substrates, including A:A mismatches and +2 insertions, showed intermediate levels of cleavage, whereas C:C and G:G pairs were cleaved with low efficiency (**Fig. 5E to G and Fig. S4**). Thus, plant AtMsh1 displays higher cleavage activity toward mismatches that commonly arise during DNA replication, such as G:T mispairs, in comparison to less frequent C:C mismatches ^44,46,47^ (**Fig. 5G and Fig. S4**). As expected, canonical G:C and A:T Watson–Crick base pairs exhibited some of the lowest levels of cleavage, with G:C cleavage being particularly weak and detectable only at higher enzyme concentrations (**Fig. 5E to G**). Although these low levels of cleavage are substantially reduced compared with those observed for mismatched substrates, the presence of these cleavage products suggests that AtMsh1 retains a degree of cleavage activity in the absence of a mismatch, particularly under conditions of elevated enzyme concentration (**Fig. 5E lanes 1 to 5 and 21 to 25 and Fig. S4**). Interestingly, despite the high binding affinity for +1 insertions and the fact that AtMsh1 function has been shown to greatly suppress the rate of 1-bp indels *in vivo* ^5,7,48^, we observed relatively low cleavage rates with the +1 and +2 insertion substrates, suggesting that the extra nucleotides may hinder proper positioning of the nuclease domain for dsDNA cleavage (**Fig. 5G and Fig. S4**).

**Fig. 5.**
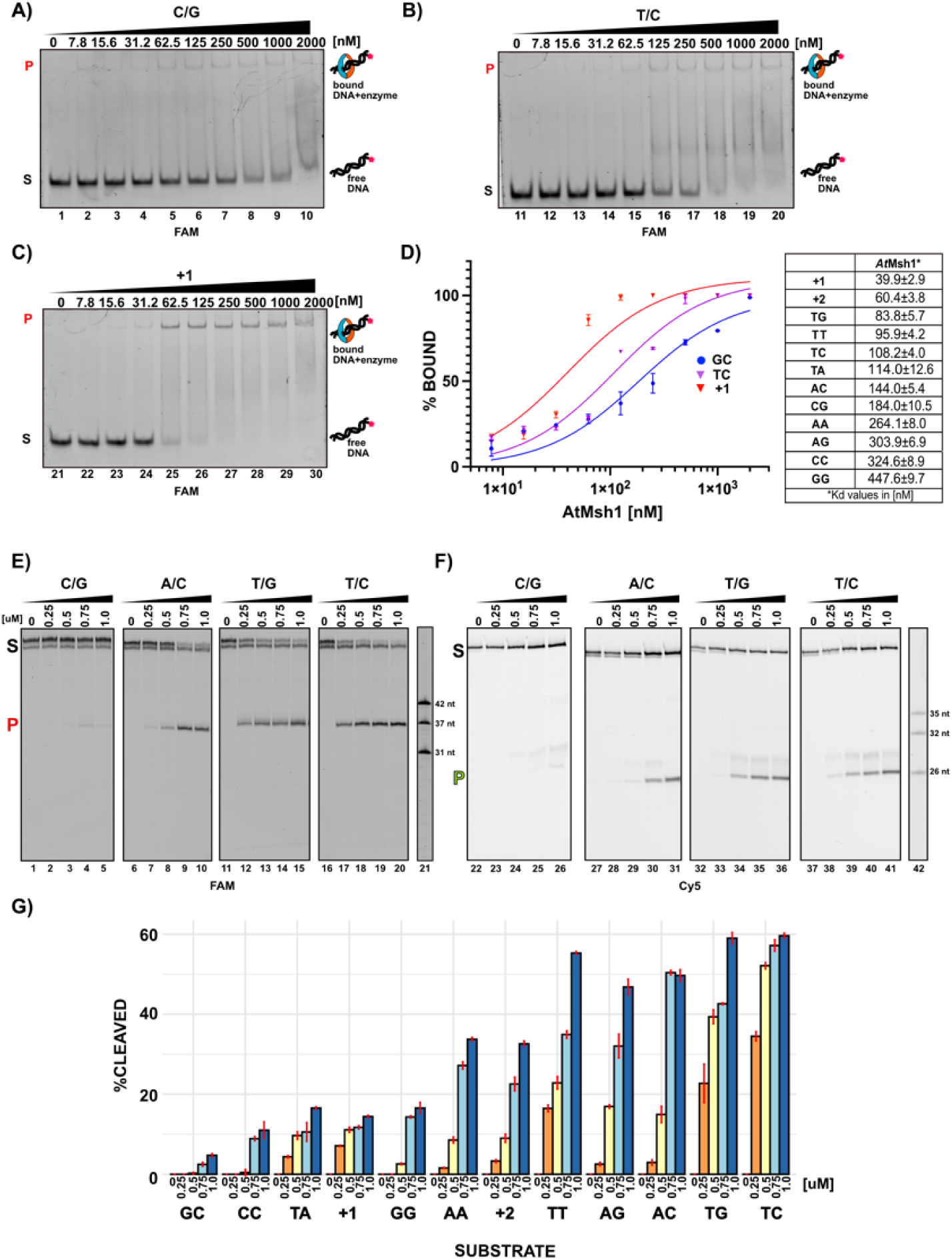
AtMsh1 exhibits preferential binding to dsDNA substrates containing indel loops and base mismatches compared with perfectly paired dsDNA and displays distinct substrate-dependent cleavage specificity. A-C) DNA binding by AtMsh1 using electrophoretic mobility shift assays (EMSA). Native gel using increasing concentrations of purified AtMsh1 incubated with fluorescently labeled DNA substrates containing a T:C mismatch, a single nucleotide indel, as well as fully complementary duplex control. **D)** Binding affinities quantified by fitting the EMSA binding data to a Langmuir binding isotherm to determine equilibrium dissociation constants (Kd) for each dsDNA substrate using data from three independent experiments. **E-F)** Cleavage activity of AtMsh1 examined using strand-specific mismatched substrates resolved on denaturing polyacrylamide gels. These assays distinguish cleavage events occurring on the mismatched strand versus the correctly paired complementary strand across selected DNA substrates, enabling evaluation of strand bias and substrate specificity in reactions incubated for 30 min at 30 °C. Oligonucleotides labeled with 5’-FAM or 5’-Cy5 are used to indicate the cleavage sites. **G)** Quantitative analysis of cleavage products to determine the percentage of substrate processed for each mismatched base pair and for 1- and 2-nt indel loops using increased AtMsh1 concentrations. Data represent the mean ± standard deviation from three independent experiments.

### AtMsh1 cleaves DNA lesions that mimic a DNA mismatch

Chemical modifications of DNA bases are a major source of mutagenesis ^49^. For example, deamination of cytosine and adenine generates uracil (U) and inosine (I), respectively, which give rise to U:A and I:C mispairs that induce C:G→T:A and A:T→G:C transitions, respectively ^49^. Oxidative damage also contributes to mutagenesis, as oxidation of guanine produces 8-oxo-7,8-dihydro-2′-deoxyguanosine (8-oxoG), a lesion that efficiently pairs with dATP promoting G:C→T:A transversions ^49^. In addition, abasic (apurinic/apyrimidinic, AP) sites are highly mutagenic, largely because DNA polymerases exhibit a strong propensity to incorporate adenine opposite them ^50–52^. To determine whether AtMsh1 exhibits substrate specificity toward additional DNA lesions, we assayed its nuclease activity on dsDNA substrates containing uracil, 8-oxoG, AP sites, or inosine. To systematically determine whether AtMsh1 preferentially cleaves damaged or mismatched DNA, we performed cleavage assays under limiting enzyme concentrations (0.25 μM) and a short incubation time (15 min). Under these conditions, AtMsh1 showed no detectable cleavage on Watson-Crick G:C or T:A pairs, whereas the T:G mismatch-containing substrate was efficiently cleaved (**Fig. 6**). Under these conditions AtMsh1 cleaves substrates containing uracil paired with either A or G (**Fig. 6A lanes 1 to 6 and 26 to 31**). Quantification of the cleavage products shows that cleavage of the U:G mismatch is more efficient than cleavage of the U:A pair and at a level comparable to a T:G mismatch (**Fig. 6B**). The intensity of the cleavage product is consistent with the nature of a U:G mismatch, as opposed to a U:A pair, in which uracil is accommodated as a canonical base and causes only minimal destabilization of the duplex due to limited disruption of base stacking ^53^ (**Fig. 6C**). Substrates containing 8-oxoG lesions paired with A or C exhibited no detectable cleavage at this AtMsh1 concentration and incubation time (**Fig. 6A, lanes 7 to 12 and 32 to 37**). This observation is consistent with the stable pairing of both the 8-oxoG:C pair and the 8-oxoG(*syn*):A mismatch, as each pair is accommodated within the dsDNA with minimal distortion and reduction in duplex stability ^54–56^. AP sites yield only barely detectable cleavage products when paired with thymine and none when paired with adenine (**Fig. 6A, lanes 14-18 and 40 to 43)**. The absence of cleavage at an abasic site is consistent with the reduced binding affinity that MutS1-family enzymes exhibit toward abasic lesions ^44^. Finally, we investigated the impact of adenine deamination using inosine-containing DNA substrates. AtMsh1 displayed selective cleavage depending on the base paired opposite inosine, showing a cleavage only when paired with Gua and Thy, and even in these cases the detected activity was much weaker than with U:G or T:G mismatches (**Fig. 6B; lanes 19 to 26 and 44 to 51)** ^57,58^. Collectively, these findings support a model in which in plant organelles AtMsh1 could function as an alternative to the base excision repair (BER) pathway, particularly in the processing of uracil-containing dsDNA, potentially providing a back-up to Uracil-DNA glycosylase activity^59,60^.

**Fig. 6.**
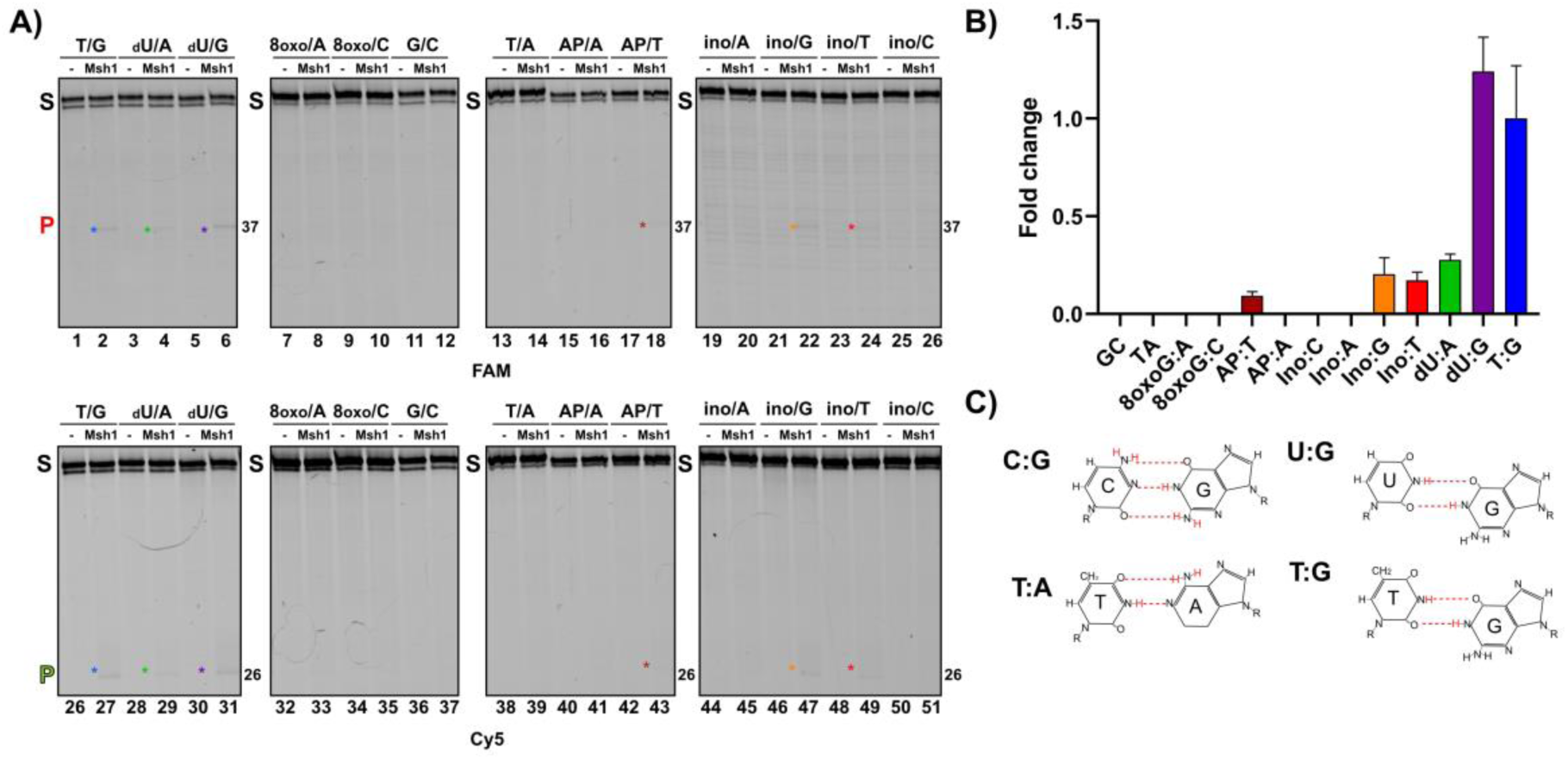
AtMsh1 cleavage across damaged DNA substrates. **A)** Denaturing polyacrylamide gel electrophoresis analysis of AtMsh1 cleavage activity on dsDNA substrates containing a T:G mismatch and Watson-Crick pairs, in comparison to uracil (U) paired with A or G, 8-oxo-guanine (8-oxoG) paired with A or C, abasic site (AP) paired with T or A, and inosine (I) paired with G,T, or C. DNA substrates were labeled with either 5′-FAM (top panels) or 5′-Cy5 (bottom panels) to monitor strand-specific cleavage. Enzymatic reactions were incubated for 15 minutes at 30 **°**C to avoid non-specific cleavage. The positions of the uncleaved substrate (S) and cleavage products (37 or 26 nts) are marked with an asterisk. **B)** Graphical representation of the fold-change in AtMsh1 cleavage activity for DNA containing lesions, relative to the T:G mismatch normalized to 1. Data represent the mean ± standard deviation from three independent experiments. **C)** Chemical structures of the DNA base pairs examined in the cleavage assays, including canonical Watson–Crick base pairs (C:G and T:A) and mismatched base pairs (U:G and T:G). Dashed red lines indicate hydrogen-bonding interactions between the paired bases.

### Divalent ions modulate AtMsh1 specificity and catalytic activity

Given that Mn²⁺ relax substrate specificity in metal-dependent enzymes and that some mismatch-specific nucleases require Mn²⁺ for activity ^28,29,61–63^, we evaluated the effect of Mn²⁺ on AtMsh1-mediated DNA cleavage using increasing concentrations of wild-type and nuclease-defective AtMsh1-Glu1078Ala (from 0.125 to 1 μM) and 5 mM of either Mg²⁺ or Mn²⁺. In the presence of Mg²⁺, AtMsh1 efficiently generated the characteristic cleavage products from the G:T mismatch substrate, with accumulation of the expected 37-nt and 26-nt fragments corresponding to cleavage of the Gua- and Thy-containing strands, respectively. Increasing enzyme concentrations enhanced the intensity of these cleavage products but did not alter the cleavage pattern or produce new cleavage products (**Fig. 7, lanes 1–5 and 21 to 25**). In the presence of Mn²⁺ and a G:T mismatch, wild-type AtMsh1 exhibits accumulation of the expected 37-nt and 26-nt fragments corresponding to cleavage of the Gua- and Thy-containing strands, respectively (**Fig. 7, lanes 6–10 and 26 to 30**) and additionally generates new cleavage products of 26, 18, and 14 nts on the Gua-containing strand and 13, 18, 37 and 43 nts on the Thy-containing strand. Reactions incubated with AtMsh1-Glu1078Ala in the presence of Mg²⁺ or Mn²⁺ did not result in a DNA cleavage product, confirming the dependence of DNA cleavage on the intact nuclease domain and the lack of contaminating nucleases during our purification protocol (**Fig. 7 lanes 11-20 and 31 to 40**). Together, these results indicate that Mn²⁺ enhances AtMsh1 catalytic activity but compromises cleavage selectivity. This suggests that the cellular balance between Mg²⁺ and Mn²⁺ concentrations may contribute to regulating the fidelity of AtMsh1-mediated DNA surveillance in plant organelles.

**Fig. 7.**
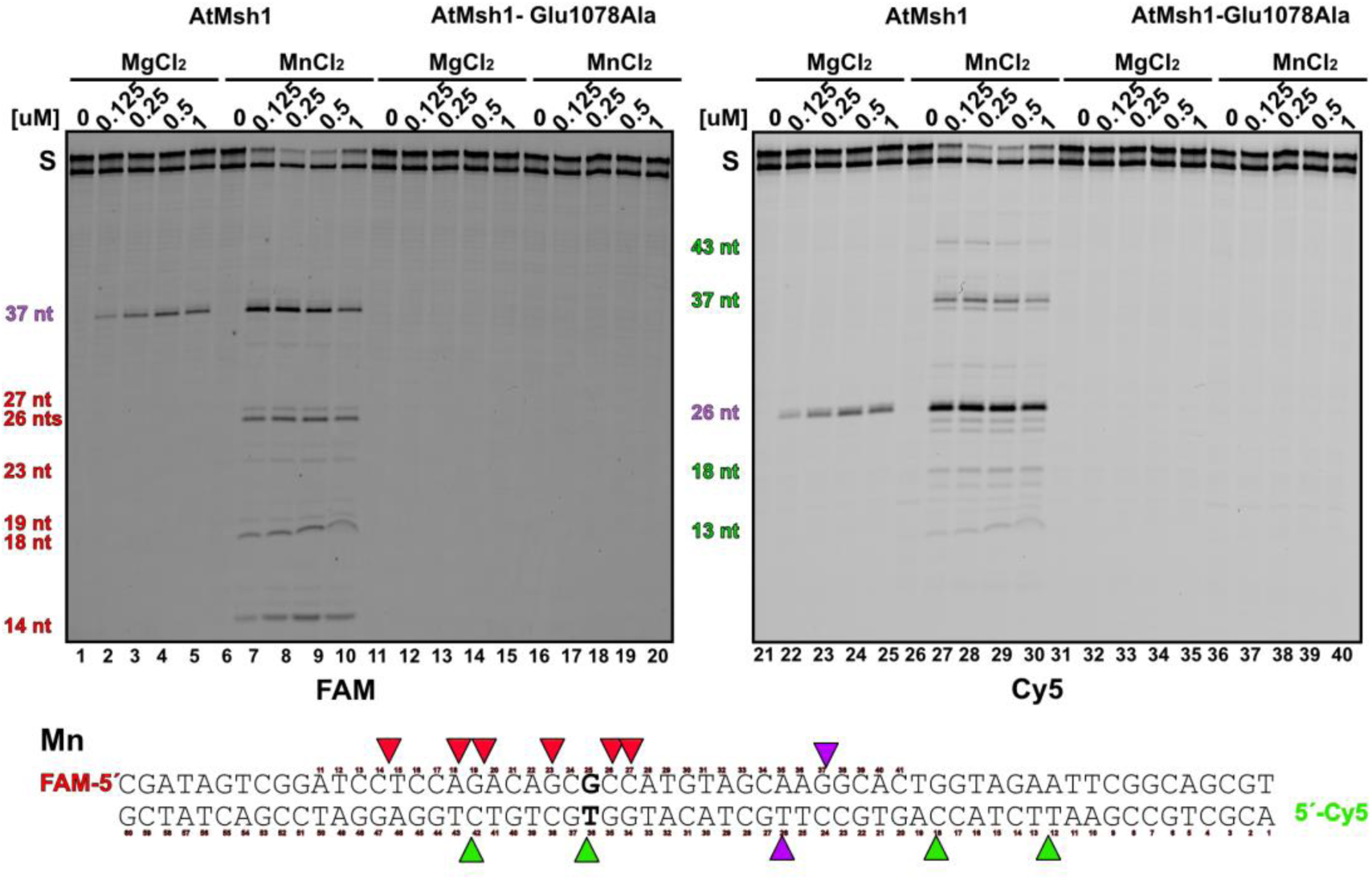
Metal-dependent cleavage of Watson-Crick and mismatch-containing dsDNA substrates by AtMsh1. 12% denaturing PAGE showing cleavage of 5′-FAM and 5′-Cy5-labeled dsDNA substrates containing a C:G dsDNA or a T:G mismatch processed by AtMsh in the presence of Mg²⁺ or Mn²⁺. In the presence of Mn²⁺, additional promiscuous double-stranded DNA cleavage products are observed for both Watson-Crick and mismatched substrates, beyond the primary cleavage site. The positions of the canonical cleavage products (P) are colored in (37 and 26 nt) are colored in purple for both strands. The promiscuous cleavage products are colored in green and red for the mismatch and complementary strands, respectively.

## DISCUSSION

Here, we have shown that AtMsh1 operates as a stand-alone mismatch-dependent endonuclease. Mechanistically, AtMsh1 recognizes DNA mismatches and lesions via its MutS1-like domain and cleaves near them via its associated GIY–YIG nuclease domain. Unlike bacterial MutS1, which preferentially recognizes and repairs T:G and A:C mismatches but exhibit reduced efficiency toward A:G, T:C, and T:T mismatches, AtMsh1 exhibits broader substrate specificity by also efficiently processing T:C, T:T, and A:G mismatches, a property that might explain the expanded mutational spectrum observed in plant organelles lacking AtMsh1 ^5,64–67^. Mitochondrial and plastid genomes in Arabidopsis *msh1* mutants are characterized by the accumulation of AT→GC and GC→AT transitions and to a lesser extent AT→CG and GC→TA transversions ^5,7^, which is consistent with the enhanced ability of AtMsh1 to recognize and cleave G:T, A:C, and T:C mismatches.

Our results also indicate that AtMsh1 recognizes and cleaves uracil embedded within dsDNA by preferentially targeting U:G mismatches and to a lesser extend U:A pairs. In contrast, AtMsh1 is not efficient in processing other DNA lesions, like 8-oxodG or abasic sites. Overall, our findings support the hypothesis that the low rate of mutation accumulation observed in plant organellar genomes results, at least in part, from an MMR-like pathway driven by the endonuclease activity of AtMsh1.^4,25,68–71^.

At present, it is unclear whether dsDNA cleavage by the GIY–YIG domain of AtMsh1 occurs through two sequential nicking reactions by a single GIY–YIG monomer, as observed in homing endonucleases, or through strand-specific nicking by opposing monomers, similar to dimeric restriction endonucleases harboring GIY–YIG domains ^17,72,73^. In comparison to GIY–YIG domains from homing endonucleases, the GIY– YIG nuclease of Msh1 enzymes harbors a predicted C-terminal helix–2–turn–helix (H2TH) motif, a common dsDNA-binding element, and a putative polyproline helix that may confer structural rigidity, resembling linker elements found in homing endonucleases (**Fig. 8**). In the presence of Mg²⁺, AtMsh1 generates unique mismatch-dependent cleavage products, introducing incisions at specific positions— nine nucleotides 5′ of the mismatch on the mismatch-containing strand and twelve nucleotides 3′ on the complementary strand in this substrate—thereby generating staggered DNA ends with three-nucleotide overhangs. This precise cleavage pattern indicates that Mg²⁺ supports a high degree of incision-site specificity. In contrast, Mn²⁺ promotes additional cleavage products at secondary sites while preserving the canonical mismatch-dependent cleavage products, suggesting that Mn²⁺ enhances catalytic activity at the expense of cleavage specificity. An important area for future research will be to identify the precise determinants of the incision site, including potential contributions local sequence context and distance from the mismatch.

**Fig. 8.**
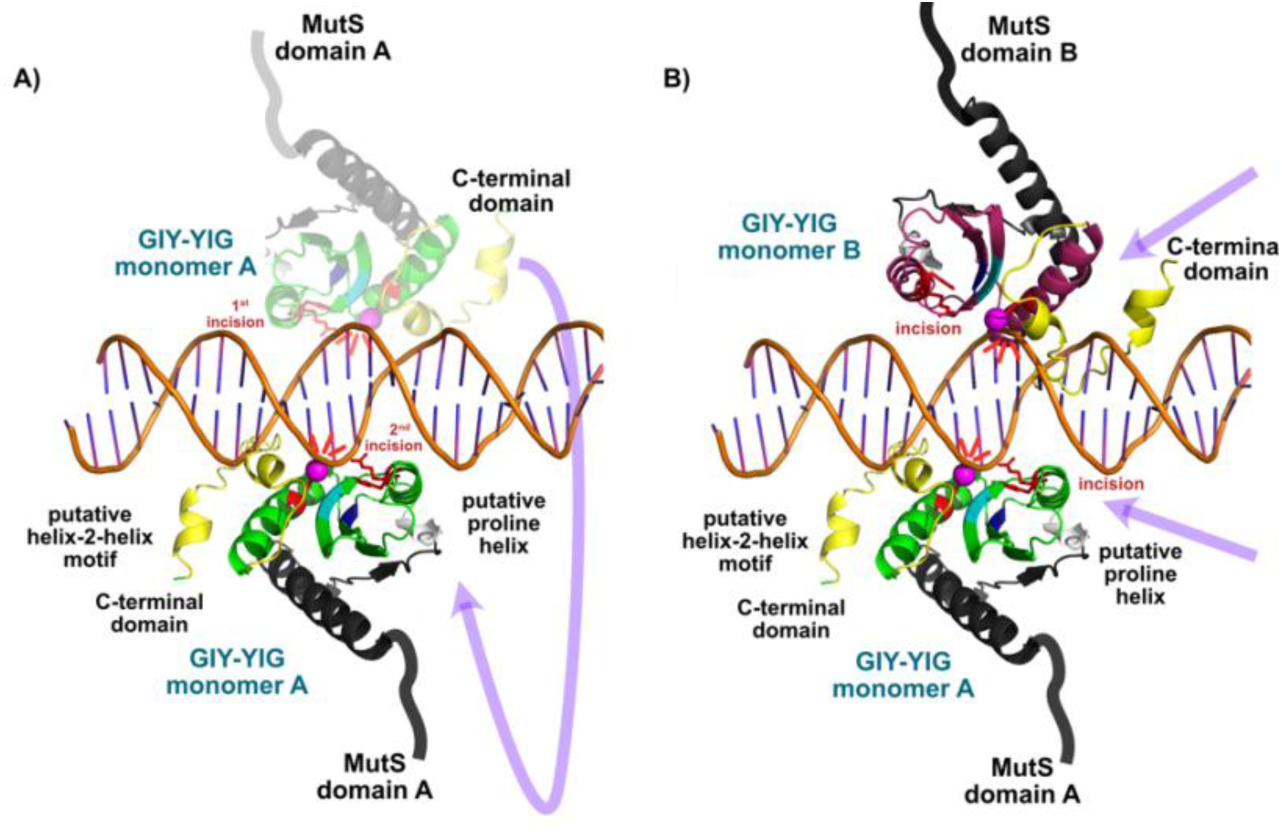
Putative models for AtMsh1 mismatch-mediated DNA cleavage. Proposed scenarios to account for the staggered nuclease cleavage of AtMsh1. (A) In the first scenario (left), a single GIY–YIG domain is “licensed” ^16^ to generate two independent incisions on opposite DNA strands, analogous to homing endonucleases that recognize a cleavage sequence and generate a first incision, and the nicked DNA reaccommodates into the nuclease active site for a second independent incision. (B) In the second model (right), each monomer acts as a nickase, contributing one incision on opposing strands, as observed for dimeric GIY–YIG restriction endonucleases. The unique structural features of the GIY–YIG nuclease of AtMsh1 (putative polyproline helix and C-terminal helix-2turn-helix (H2TH) motif are colored in gray and yellow, respectively). The metal ion that coordinates phosphodiester DNA incisions are colored in magenta, and the conserved catalytic triad (Arg1048, His1052, and Glu1078) are represented in a stick representation and colored in red. The dsDNA is modeled based on a superimposition of the AtMsh1 domain on the GIY–YIG nuclease from the crystal structure of the restriction enzyme Eco29kI^17^.

Elucidating if AtMsh1 distinguishes between newly incorporated errors is essential for understanding organellar MMR in plant organelles ^74,75^. Unlike EndoMS/NucS nucleases, which excise dsDNA sections containing the mismatched or lesion-containing bases ^76–78^, AtMsh1 introduces incisions several nucleotides from the mismatch in our substrates generating staggered DNA ends. We propose that these distal cleavage events generate entry sites for 5′→3′ and 3′→5′ plant organellar exonucleases (e.g. OEX 1 and 2, and organellar exonuclease DPD1, respectively ^79,80^), which may remove the mismatch or lesion and generate 3′ single-stranded overhangs suitable for plant organellar RecA-mediated homologous recombination. Consistent with this model, plant mitochondria and plastids harbor enzymes required for homologous recombination and contain multiple copies of organellar genomes ^81–84^. These findings are consistent with the antimutagenic role of AtMsh1, as demonstrated *in vivo* using mitochondrial TALE–cytidine deaminases in Arabidopsis^85^, and support a model in which the exceptionally low mutation rates of plant organellar genomes are maintained by a unique DNA repair pathway in which AtMsh1 mismatch-directed dsDNA cleavage is followed by exonucleolytic resection and homologous recombination, ultimately promoting gene conversion.

## ACKNOWLEDGMENTS

To Corina Díaz-Quezada for technical assistance and Dr. Antolín Peralta and Dr. Rachael DeTar for helpful discussions. APA thanks CONACYT(SECIHTI)-Mexico for his graduate student fellowship. NBT is a Pew Latin American Postdoctoral Fellow.

## AUTHOR CONTRIBUTIONS

**APA and CZ:** Investigation, formal analysis, validation, visualization, writing. **NBT:** Formal analysis, investigation. **DBS:** Formal analysis, writing – review & editing. **SA and LGB:** Conceptualization, investigation, funding acquisition, supervision, formal analysis, writing – review & editing.

## SUPPLEMENTARY DATA

Supplementary Data are available online.

## CONFLICT OF INTEREST

The authors declare no conflict of interest.

### Data availability statement

All data are incorporated into the article and its online supplementary material

