## Supplemental information for "Plant MutS Homolog 1 is a mismatch-directed nuclease required for organelle genome maintenance"

Supplementary Information

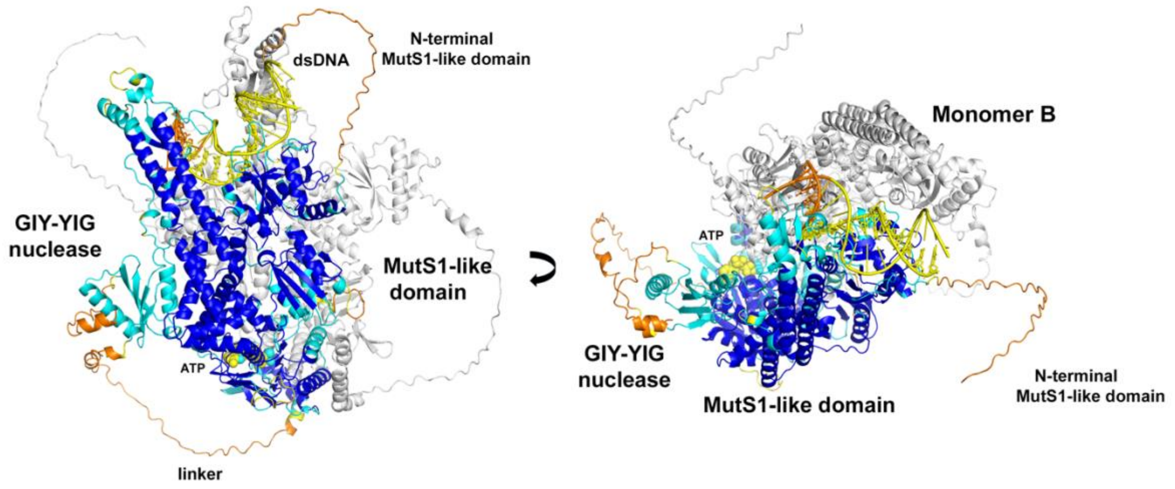


**Fig. S1 AlphaFold 3 model of AtMsh1 with based on Taq MutS1 bound to T:G mismatch.** AtMsh1 is modeled as a homodimer, with the mismatch contacting monomer colored according to pLDDT values and the second monomer shown in white. pLDDT values indicate the confidence of the predicted AtMsh1 structure, with blue (>90) representing very high confidence, light blue/cyan (70–90) reliable predictions, yellow (50–70) low confidence, and orange/red (<50) very low confidence. The modeled ATP and catalytic metal ion bound to the ATPase domain are represented as spheres. The AtMsh1 model shows the conserved overall architecture of MutS1 proteins, together with an additional C-terminal GIY–YIG nuclease domain.

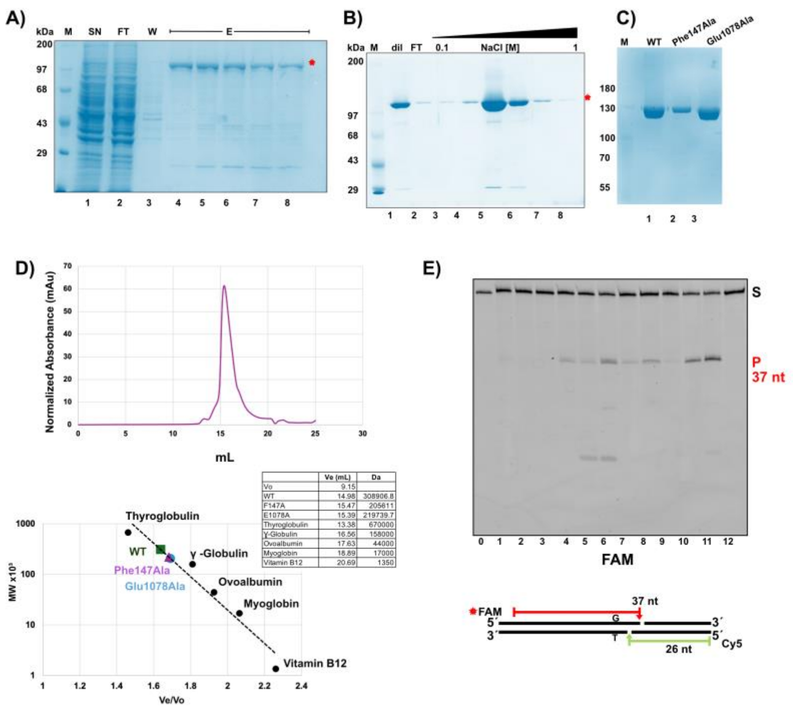


**Fig. S2 Recombinant purification of yeast-expressed AtMsh1 and identification of optimal buffer conditions for nuclease activity. A and B**) IMAC and Heparin affinity chromatography profile of yeast-purified AtMSH1, with fractions analyzed by 10% SDS–PAGE and Coomassie Blue staining. The analysis reveals a single major purified protein with a high degree following those steps. **C)** Wild-type and point mutants of AtMsh1 obtained after size-exclusion chromatography run on a 10% SDS–PAGE gel and detected by Coomassie blue staining. **D)** Size-exclusion chromatography oof wild-type AtMSh1 on a Sephacryl S-300 column, calibrated with molecular-weight standards, showed that AtMsh1 eluted as a single major peak at a volume consistent with a dimeric AtMsh1 complex. Peak fractions were analyzed by 10% SDS–PAGE followed by Coomassie Blue staining, confirming the presence of a highly purified protein species. A minor peak eluting in fractions 19 to 21 corresponded to contaminant proteins 1. **E) Identification of optimal reaction conditions for AtMsh1 nuclease activity.** Representative 12% denaturing polyacrylamide gel showing cleavage of a 5′-FAM-labeled T:G mismatch-containing DNA substrate by yeast-purified AtMsh1 in twelve reaction buffers. Efficient mismatch-dependent cleavage was observed under several conditions, with maximal activity in buffer 11, which was used in all subsequent experiments. The positions of the uncleaved substrate (S) and cleavage products (P) are indicated. Buffer compositions were as follows: **(1)** NEB1: 10 mM Bis-Tris propane-HCl (pH 7.0), 10 mM $MgCl_2$, 1 mM DTT; **(2)** NEB2: 10 mM Tris-HCl (pH 7.9), 10 mM $MgCl_2$, 50 mM NaCl, 1 mM DTT; **(3)** NEB3: 50 mM Tris-HCl (pH 7.9), 100 mM NaCl, 10 mM $MgCl_2$; **(4)** NEB4: 20 mM Tris-acetate (pH 7.9), 50 mM potassium acetate, 10 mM magnesium acetate; **(5)** Thermo Amplification Grade DNase I buffer: 200 mM Tris-HCl (pH 8.4), 20 mM $MgCl_2$, 500 mM KCl; **(6)** CutSmart buffer: 20 mM Tris-acetate (pH 7.9), 50 mM potassium acetate, 10 mM magnesium acetate; **(7)** R buffer (Modified MutY buffer): 40 mM Tris-HCl (pH 7.5), 150 mM NaCl, 5 mM $MgCl_2$; **(8)** F buffer (modified from Fukui *and cowokers* [34]*):* 50 mM Tris-HCl (pH 7.5), 100 mM KCl, 5 mM $MgCl_2$, 1 mM DTT; **(9)** N buffer (modified NucS buffer): 25 mM Bis-Tris (pH 6.4), 2.5 mM $MgCl_2$, 5 mM DTT, 0.1% Triton X-100; **(10)** BTP 6.5: 25 mM Bis-Tris propane (pH 6.5), 50 mM potassium glutamate, 1 mM DTT, 5 mM $MgCl_2$; **(11)** BTP 7.1: 25 mM Bis-Tris propane (pH 7.1), 50 mM potassium glutamate, 1 mM DTT, 5 mM $MgCl_2$; and **(12)** Original buffer assayed in Peñafiel-Ayala and coworkers [14]: 50 mM Tris-HCl (pH 8.0), 150 mM NaCl, 5 mM $MgCl_2$, 1 mM DTT. All reactions were supplemented with 0.1 mg/mL bovine serum albumin (BSA) and 2 mM ATP.

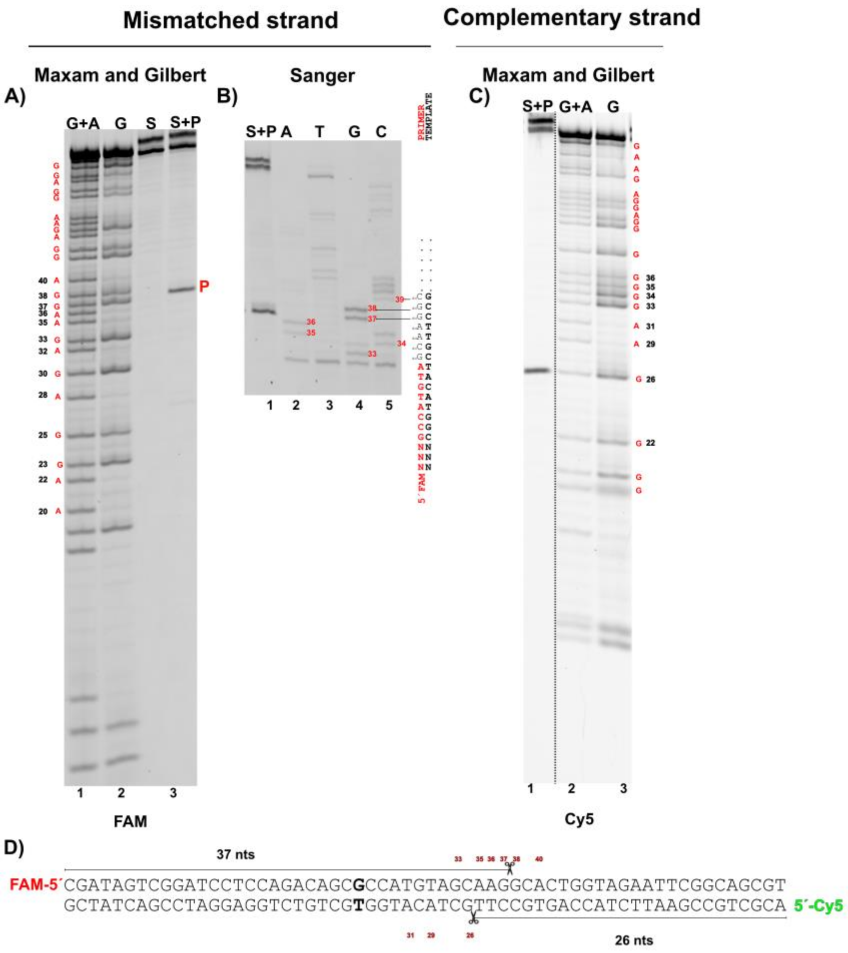


**Fig. S3. Mapping of the AtMsh1 cleavage site by Maxam–Gilbert and Sanger sequencing.** The cleavage site of AtMsh1 was determined by comparison of the cleavage product with sequencing ladders generated by Maxam–Gilbert cleavage and Sanger dideoxy sequencing. **A)** Maxam-Gilbert reactions on the 5′-FAM-labeled strand matches with the destruction between the phosphodiester bond at bases 37 and 38. **B**) Sanger sequencing places the cleavage site at the ddGTP incorporated at position 37 from the 5´-end labeled oligonucleotides, supporting that cleavage at the FAM-labeled strand proceeds between bases 37 and 38. **C)** The cleavage product of the 5′-Cy5-labeled strand containing strand matches the hydrolytic degradation of Gua at position 27 from the 5´-end labeled oligonucleotide. As the chemical modification in a Maxam-Gilbert reaction induces the destruction of the base, the cleavage site is located between nucleotides 27 and 26. Thus, the analysis indicates that AtMsh1 generates a 5′-FAM-labeled 34-nt dsDNA fragment with a 3-nt 3′ overhang, along with a 5′-Cy5–labeled 23-nt dsDNA fragment that also carries a 3-nt 3′ overhang. **D)** Analysis of cleavage products generated from the mismatched DNA substrate. The labeled strand and predicted cleavage sites are indicated in color and by scissors, respectively.

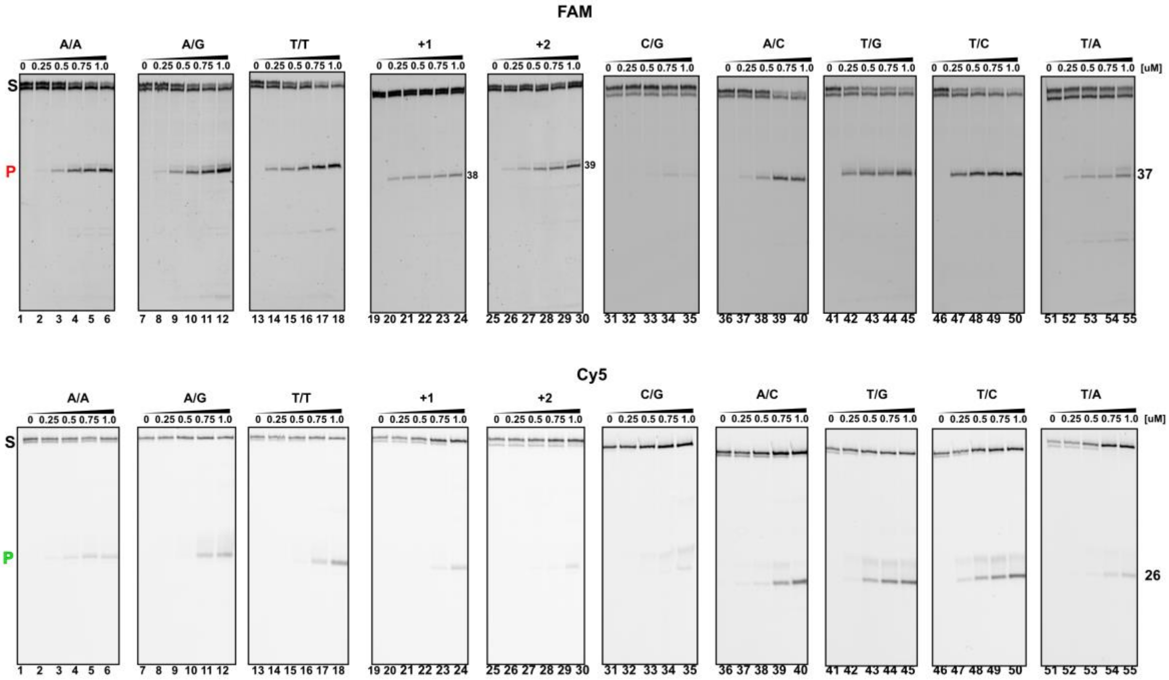


**Fig. S4 Cleavage activity of AtMsh1 on all possible DNA mismatches and +1/+2 nucleotide indels.** Denaturing 12% polyacrylamide gel electrophoresis showing cleavage of 5′-FAM and 5′-Cy5-labeled double-stranded DNA substrates by AtMsh1. Upper panels show cleavage of substrates labeled on the 5′-FAM DNA strand, and lower panels show cleavage of the 5′-Cy5 labeled complementary strand. Substrates contained all possible single-base mismatches (A:A, A:C, T:G, T:C, G:G, C:C, A:G, T:T, G:A, C:T, and G:C and A:T as Watson–Crick controls) or indel loops of +1 and +2 nucleotides. Cleavage reactions were performed in the presence of $Mg^{2+}$. The positions of the uncleaved substrate (S) and the characteristic cleavage products (P) are indicated.

### Table S1. Oligonucleotides used for AtMsh1 cloning and site-directed mutagenesis.

| NAME | SEQUENCE 5´-3´ |
|---|---|
| Glu1078Ala-FWD | ATGTCAATTGgcaACATTATTAATCAAC |
| Glu1078Ala-REV | GCCATGGATTTACCTTGC |
| Phe147Ala-FWD | AGTTGGTGAAgctTACGAAGCAATTG |
| Phe147Ala-REV | CTACACAATAAAACTTCTCTTG |
| pSc-Msh1-BamHI | GAAGAGGGATCCGTAACCATGTCACACCATCACCATCACCATCACCATCACGGCAGCAGCGGCATGCCATCTTCTCCAATTTTATTGAACAGA |
| Msh1-XhoI | CTCGATTCTCGAGTTACAAAATAGAAACAACATCT |

**Table S2. Oligonucleotide sequences used in this study to assemble DNA substrates containing different mismatches and lesions and to generate molecular rulers.**

| Oligonucleotide | SEQUENCE |
|---|---|
| Ladder26cy5 | Cy5 5´ACGCTGCCGAATTCTACCAGTGCCTT |
| Ladder42FAM | 6-FAM 5´CGATAGTCGGATCCTCCAGACAGCGCCATGTAGCAAGGCACT |
| Ladder37FAM | 6-FAM 5´CGATAGTCGGATCCTCCAGACAGCGCCATGTAGCAAG |
| Ladder31FAM | 6-FAM 5´CGATAGTCGGATCCTCCAGACAGCGCCATGT |

| | |
|---|---|
| 25_Oxo-Dan | 6-FAM 5´CGATAGTCGGATCCTCCAGACAGC/oxoG/CCATGTAGCAAGGCACTGGTAGAATTCGGCAGCGT 3´ |
| 25_dU-Dan | 6-FAM 5´CGATAGTCGGATCCTCCAGACAGC/dU/CCATGTAGCAAGGCACTGGTAGAATTCGGCAGCGT 3´ |
| 25_ino-Dan | 6-FAM 5´CGATAGTCGGATCCTCCAGACAGC/In/CCATGTAGCAAGGCACTGGTAGAATTCGGCAGCGT |
| 25_AP-Dan | 6-FAM 5´CGATAGTCGGATCCTCCAGACAGC/idSp/CCATGTAGCAAGGCACTGGTAGAATTCGGCAGCGT 3´ |
| PolEsc32_Cy5 | Cy5 5´ACGCTGCCGAATTCTACCAGTGCCTTGCTACA 3´ |
| PolEsc35_Cy5 | Cy5 5´ACGCTGCCGAATTCTACCAGTGCCTTGCTACATGG 3´ |

| | |
|---|---|
| FAM_Thy | 6-FAM 5´CGATAGTCGGATCCTCCAGACAGCTCCATGTAGCAAGGCACTGGTAGAATTCGGCAGCGT 3´ |
| FAM_Ade | 6-FAM 5´CGATAGTCGGATCCTCCAGACAGCACCATGTAGCAAGGCACTGGTAGAATTCGGCAGCGT 3´ |
| FAM_Gua | 6-FAM 5´CGATAGTCGGATCCTCCAGACAGCGCCATGTAGCAAGGCACTGGTAGAATTCGGCAGCGT 3´ |
| FAM_Cyt | 6-FAM 5´CGATAGTCGGATCCTCCAGACAGCCCCATGTAGCAAGGCACTGGTAGAATTCGGCAGCGT 3´ |
| Cy5_Gua | Cy5 5´ACGCTGCCGAATTCTACCAGTGCCTTGCTACATGGGGCTGTCTGGAGGATCCGACTATCG 3´ |
| Cy5_Cyt | Cy5 5´ACGCTGCCGAATTCTACCAGTGCCTTGCTACATGGCGCTGTCTGGAGGATCCGACTATCG 3´ |
| Cy5_Ade | Cy5 5´ACGCTGCCGAATTCTACCAGTGCCTTGCTACATGGAGCTGTCTGGAGGATCCGACTATCG 3´ |
| Cy5_Thy | Cy5 5´ACGCTGCCGAATTCTACCAGTGCCTTGCTACATGGTGCTGTCTGGAGGATCCGACTATCG 3´ |
| T_ins1 | 6-FAM 5´CGATAGTCGGATCCTCCAGACAGCTTCCATGTAGCAAGGCACTGGTA*GAATTC*GGCAGCGT 3´ |
| T_ins2 | 6-FAM 5´CGATAGTCGGATCCTCCAGACAGCTTTCCATGTAGCAAGGCACTGGTA*GAATTC*GGCAGCGT 3´ |

**Table S3. Oligonucleotide combinations used to generate double-stranded DNA substrates**

| substrate | OLIGONUCLEOTIDE COMBINATION |
|---|---|
| T/A | Cy5Thy+FamAde |
| T/G | Cy5Thy+FamGua |
| T/C | Cy5Thy+FamCyt |
| T/T | Cy5Thy+FamThy |
| A/C | Cy5Ade+FamCyt |
| A/G | Cy5Ade+FamGua |
| A/A | Cy5Ade+FamAde |
| G/G | Cy5Gua+FamGua |
| C/C | Cy5Cyt+FamCyt |
| C/G | Cy5Gua+FamCyt |
| C/A | Cy5Ade+FamCyt |
| ins_1 | Cy5Ade+T_ins1 |
| ins_2 | Cy5Ade+T_ins2 |
| 8oxoG/A | 25_Oxo + Cy5Ade |
| 8oxoG/C | 25_Oxo + Cy5Cyt |
| AP/T | 25_AP + Cy5Thy |
| AP/A | 25_AP + Cy5Ade |
| dU/G | 25_dU + Cy5Gua |
| dU/A | 25_dU + Cy5Ade |
| Ino/A | 25_ino + Cy5Ade |
| Ino/G | 25_ino + Cy5Gua |
| Ino/T | 25_ino + Cy5Thy |
| Ino/C | 25_ino + Cy5Cyt |

4
